# Comparison of Mitogenomic and Mitochondrial Markers for the Phylogenetic and Evolutionary Study of Marmosets, with special focus on captive *Callithrix jacchus*

**DOI:** 10.1101/2022.09.19.508604

**Authors:** João L.G. Fonseca, Ricardo C. H. del Rosario, Maria Adélia Borstelmann de Oliveira, Guoping Feng, Claudia S. Igayara, Steven A. McCarroll, Patricia A. Nicola, Luiz C. M. Pereira, Jeffery Rogers, Christian Roos, Joanna Malukiewicz

## Abstract

Marmosets, especially *Callithrix jacchus*, are becoming valuable, high-demand biomedical models but their supply outside of their native Brazil is limited. Thus, uncovering the ancestry and biogeographic origins of captive marmosets is an essential task to optimize their biomedical use. Such information facilitates assessing standing levels of captive marmoset genetic diversity, identifying hybrids, and maintaining genetic diversity of captive marmoset populations. Here, we present 16 newly sequenced mitochondrial genomes from marmosets with *C. jacchus* phenotypes from international biomedical and animal supply facilities. We combine these new data with publicly available sequences to compare usage of mitogenome and the mtDNA control region for determining the ancestry and diversity of captive marmosets. MtDNA control region and mitogenome haplotypes from all 16 newly sequenced marmosets grouped within *C. jacchus* phylogenetic clades. Overall, phylogenies based on mitogenomes rather than the control region showed better resolved branching patterns with stronger statistical support within *C. jacchus* clades. However, both mtDNA markers show that *C. jacchus* is the sister clade to the phylogenetic *C. penicillata* Caatinga biome clade. *Callithrix jacchus* mtDNA control region haplotypes showed the highest haplotype and nucleotide diversity, and were intermediate in terms of per sequence theta and polymorphic sites. The larger number of available *Callithrix* mtDNA control region sequences relative to mitogenomes was used to determine ancestral mtDNA haplotype biogeographic origins. However, biogeographic origins for many *C. jacchus* haplotypes including those from international captive facilities could not be determined, which we attributed to unknown provenance information from natural *C. jacchus* populations. Given our results, we encourage increased sampling and genomic sequencing efforts for natural *C. jacchus* populations in Brazil, and utilizing the mitogenome over other mtDNA markers for phylogenetic-based analyses. Further, combining marmoset mitogenomic data with phenotypic and nuclear genomic data will greatly enhance the biomedical use of marmosets.

## 1. Introduction

*Callithrix* marmoset species, which are endemic to Brazil (Figure 1), possess a unique set of biological characteristics such as frequent twinning and rapid reproductive rate [1]. Such traits have resulted in marmosets becoming valuable biomedical models that are in high demand among international research facilities, but whose supply is limited [2]. It is estimated that the worldwide population of marmoset biomedical colonies comprises approximately 6,000 marmosets, of which 2,500 are found in Japan, 1,900 in North America, 1,000 in Europe, and 800 in South America [2]. Within the US, marmosets are housed across approximately 27 research facilities, and the founder marmosets imported into the US during the 1960s and 1970s came from Europe and the United Kingdom [2]. Pedigree information on the marmosets initially imported into the US is limited, but in many cases present-day marmoset colonies originate from stocks of a few dozen imported animals [2].

**Figure 1:**
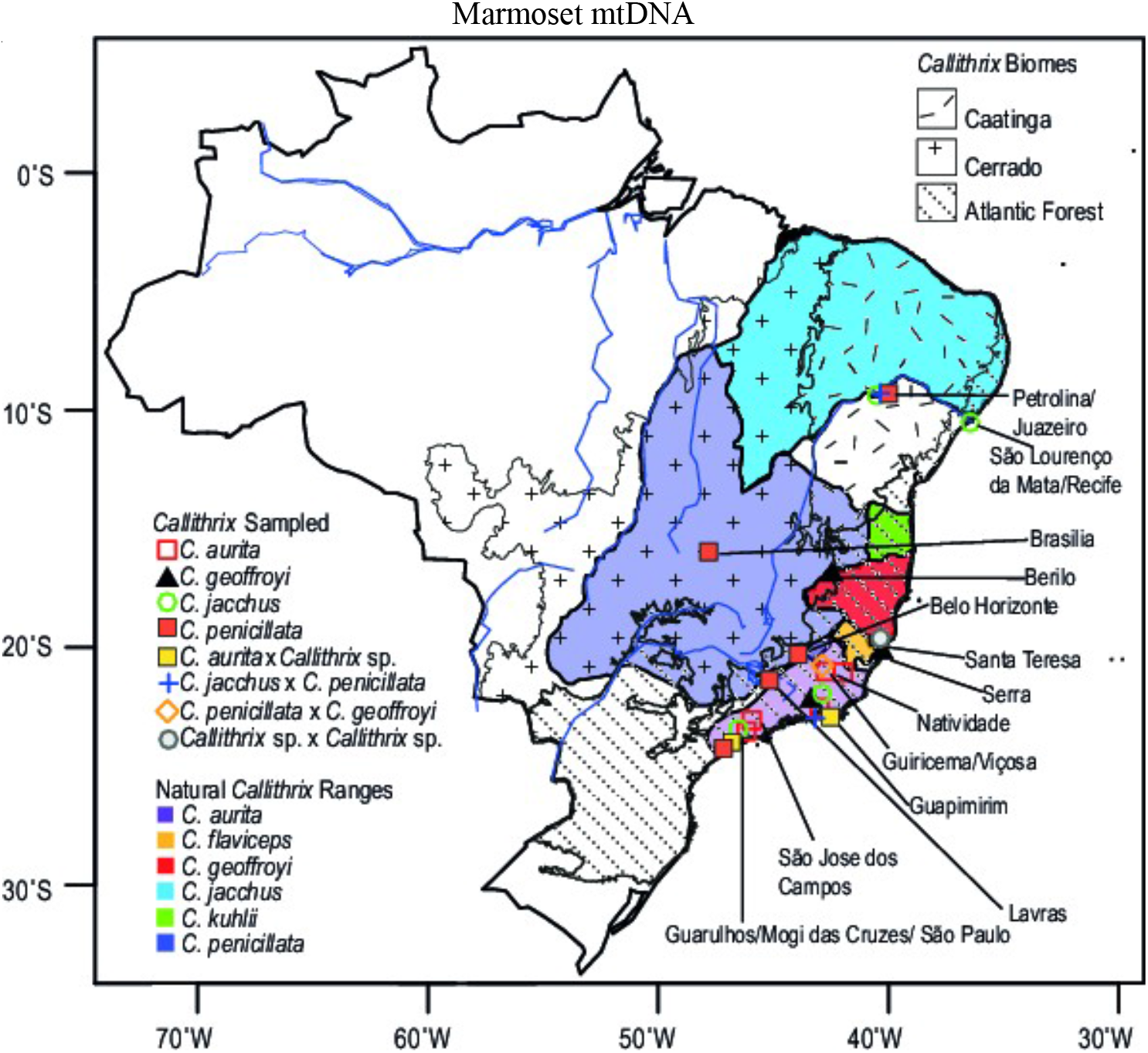
Approximate distribution of *Callithrix* species in Brazil (2012 IUCN Red List Spatial Data; (http://www.iucnredlist.org/technical-documents/spatial-data) and geographic origins of mtDNA control region sequences available from previously published studies. Locations of three biomes where *Callithrix* occur naturally, the Caatinga, Cerrado, and Atlantic Forest, are also indicated.

Research facilities generally maintain colonies that consist of individuals with the phenotype of common marmosets (*C. jacchus*). This species is characterized by white bushy ear tufts and a white forehead star, but the ancestral biogeographic origins of these captive monkeys are largely unknown. In Brazil, common marmosets occur within three major Brazilian biomes: the scrub-brush and semi-arid Caatinga, the semiarid savanna-like Cerrado, and the humid-coastal Atlantic Forest [1, 3]. Marmosets are also known to hybridize, both under free-ranging and captive conditions, with the majority of cases involving *C. jacchus* as one of the parental species [1, 4–6]. Brazilian free-ranging populations of common marmosets are declining [7], and marmosets for biomedical use are difficult to import under international wildlife trade regulations [2]. Given the limited marmoset supply outside of Brazil, uncovering the phylogeny, ancestral biogeographic origins, and hybrid status of captive marmosets is critical for their biomedical use. Such information is helpful in assessing standing levels of captive marmoset genetic diversity, identifying possible hybrids, and for maintaining genetic integrity of captive, biomedical marmoset populations.

Mitochondrial DNA (mtDNA) has been a valuable and informative genetic marker in resolving *Callithrix* phylogeny, divergence, biogeography, and hybrid identification [4, 6, 8–10]. Patterns of interspecific divergence analyzed using the approximately 1200 base pair (bp) rapidly-evolving mtDNA control region, which is much smaller than the approximately 16,500 bp *Callithrix* mitogenome, are largely concordant with results from full *Callithrix* mitogenome analyses [4, 6, 8–10]. Here we report on a comparison of the mitogenome and the mtDNA control region for determining the ancestry and diversity of marmosets from international research and animal supply facilities by combining newly and previously available sequence data. We utilized both data types to address the following questions: (1) What are the ancestral origins of marmosets maintained at international captive facilities?; (2) Are cryptic hybrids present among international captive marmoset populations?; (3) How does the genetic diversity of *C. jacchus* mtDNA control region and mitogenome compare to that of other *Callithrix* taxa?

## 2. Methods

For this study, we newly sampled genetic material from a total of 16 marmosets with a *C. jacchus* phenotype from the New England Primate Research Center (NEPRC), a US animal supplier (Company A) and one Japanese animal supplier (CLEA, Tokyo, Japan). We also sequenced the mitogenome of a captive marmoset (BJT23) with a *C. aurita* phenotype whose skin sample was collected in 2015 in Brazil. Furthermore, we obtained publicly available and previously published mtDNA control region and mitogenomic sequences from GenBank from a total of 266 marmosets from five different *Callithrix* species and 4 different hybrid types (Table 1). Detailed information on marmoset mitogenomic and mtDNA control region sequences is given in Supplementary Table S1.

**Table 1.** Sampling Summary of *Callithrix*

| Classification | Count |
| --- | --- |
| *C. aurita* | 13 |
| *C. geoffroyi* | 16 |
| *C. jacchus* | 103 |
| *C. kuhlii* | 9 |
| *C. penicillata* | 55 |
| *C. jacchus* x *C. penicillata* | 53 |
| *C. penicillata* x *C. geoffroyi* | 11 |
| *C. aurita* x *Callithrix* sp. | 5 |
| *Callithrix* sp. x *Callithrix* sp. | 1 |
| Total | 226 |

Captive marmosets outside of Brazil were collected under the approval of the Massachusetts Institute of Technology IACUC (protocol number 051705020). Tissue from BJT23 was collected under the approval of the ASU Institutional Animal Care and Use Committee Animals (ASU IACUC, protocols #11-1150R, 15-144R) and Brazilian Environmental Ministry (SISBIO protocols #47964–2 and #28075–2). The Brazilian marmoset sample BJT23 was registered in the Brazilian SISGEN database under number A2E885E. Biological tissue sampling complied with all institutional, national, and international guidelines. Species phenotypic classifications of newly sampled marmosets were based on published phenotypic descriptions following [1].

### 2.1 Laboratory and Sequencing Protocol

To obtain the control region sequence of individual BJT23, we sequenced the entire mitogenome following the long-range PCR strategy of [6, 10]. To check for the presence of nuclear copies of mitochondrial genes (numts) in the mitogenone sequence of BJT23, we first checked the sequence electropherogram which showed high quality base calls with no ambiguous nucleotides, indicating absence of different populations of mitochondrial-like sequences [10]. Then, sequences of all 13 mtDNA protein-coding genes were translated into protein sequences to check for premature stop codons and frame shifting insertions or deletions [10, 11]. Next, neighbor-joining trees were con-structed with MEGA 11 [12] for each protein-coding gene using BJT23 sequences and those from previously available mitogenome sequences described above to check for unusual phylogenetic placement of species sequences. These analyses did not reveal any unexpected amino acid translation or phylogenetic patterns, therefore mitogenomic sequences from BJT23 were assumed to be free of numts.

For remaining samples, an aliquot of genomic DNA (150ng in 50*µ*L) was used as the input for DNA fragmentation. Fragmentation was performed using a Covaris focused-ultrasonicator, targeting 385bp fragments. Following fragmentation, additional size selection was performed using a SPRI cleanup. Library preparation was performed using a commercially available kit provided by KAPA Biosystems (KAPA Hyper Prep with Library Amplification Primer Mix, product KK8504),with palindromic forked adapters using unique 8-base index sequences embedded within the adapter (purchased from Roche). The libraries were then amplified by 10 cycles of PCR. Following sample preparation, libraries were quantified using quantitative PCR (kit purchased from KAPA Biosystems) with probes specific to the ends of the adapters. This assay was automated using Agilent’s Bravo liquid handling platform. Based on qPCR quantification, libraries were normalized to 2.2nM and pooled into 24-plex libraries.

The 24-plex sample pools were combined with NovaSeq Cluster Amp Reagents DPX1, DPX2 and DPX3 and loaded into single lanes of a NovaSeq 6000 S4 flowcell using the Hamilton Starlet Liquid Handling system. Cluster amplification and sequencing were performed on NovaSeq 6000 Instruments utilizing sequencing-by-synthesis kits to produce 151 bp paired-end reads. Output from Illumina software was processed by the Picard data-processing pipeline to yield CRAM or BAM files containing demultiplexed, aggregated aligned reads.

### 2.2 Mitogenome and mtDNA Control Region Assembly from Whole Genome Sequences

For samples that were included in the Illumina sequencing protocol described above, all newly sequenced *Callithrix* mitogenomes were assembled from whole genome data following a modified protocol of [6]. Our modifications included using the bwa-mem algorithm of BWA 0.7.12 [13] to align Illumina short-reads from individual marmoset whole genome data to a composite reference genome comprised of the *C. jacchus* ASM275486v1 genomic assembly and *C. jacchus* mitochondrial genome (GenBank Accession# NC_025586.1). For each marmoset sample, reads that aligned to the mitochondrial genome were gathered with SAMTOOLS 1.8 using the command “samtools view -h -f 3 -F 2316 NC_025586.1 $BAM | samtools collate -Ou - | samtools bam2fq -t -1 $READ1 -2 $READ2 –2.” Then mitogenomes were each assembled with NOVO-PLASTY 2.6.4 [14], for which config files were created for each individual marmoset that included the path of mitogenome-aligned forward and reverse fastqs (\$READ1 and \$READ2) from the previous reference genome mapping step. Subsequently, mtDNA control region sequences were manually extracted from these new mitogenomes after annotation following the protocol of [6]. All newly sequenced mtDNA control region and mitogenome sequences have been deposited in GenBank (Table S1). We also extracted mtDNA control region sequences from *Callithrix* mitogenome sequences downloaded from GenBank.

### 2.3 mtDNA Control Region Alignment and Data Analysis

All mtDNA control region sequences were aligned in MAAFT (https://www.ebi.ac.uk/Tools/msa/mafft/) with default settings and MAFFT alignments were confirmed visually in Mesquite 3.5 [15]. The initial alignment of 1125 bases was trimmed to 881 bases to accommodate for the length of all used sequences. Haplotypes for our dataset were determined with DnaSP 6.12.03 [16], and haplotypes were used for phylogenetic and network reconstruction. The mitogenome and control region sequences of *Cebuella pygmaea* (GenBank accession #MZ747454.1) were used as a phylogenetic outgroup. We reconstructed phylogenetic trees with maximum likelihood (ML) and Bayesian algorithms using IQ-TREE 1.6.12 [17] and MrBayes 3.2.7 [18, 19]. For the ML phylogeny, we used the optimal substitution model “HKY+F+I+G4” as calculated with ModelFinder [20, 21] in IQ-TREE under BIC. We performed the ML analysis two separate times in IQ-TREE with 1000 ultrafast bootstrap (BS) replications [22] per run. The two resulting ML trees were then compared visually for concordance. For the Bayesian tree, we used Modeltest-ng 0.1.6 [23] with the -T flag set to “mrbayes” to determine the optimal substitution model as “HKY + I + G4” under BIC. The Bayesian tree was reconstructed via Markov Chain Monte Carlo (MCMC) runs with 3,000,000 generations and tree and parameter sampling occurring every 100 generations. Upon completion of the two runs, the first 25% of generations were discarded as burn-in. To check convergence of all parameters and the adequacy of the burn-in, we assessed the uncorrected potential scale reduction factor (PSRF) [24] and that all parameter Estimated Sample Size (ESS) values were above 200. Phylogenetic trees were visualized and edited with FigTree 1.4.2 (http://tree.bio.ed.ac.uk/software/figtree/). Designations of resulting ML and Bayesian phylogenetic clades followed that of [6].

Due to the large number of *Callithrix* control region sequences available with known sample provenance, we took advantage of this information to reconstruct the biogeographic history of *Callithrix* mtDNA control lineages. Using a user supplied phylogenetic tree, ancestral *Callithrix* geographic ranges were obtained with the methodology of [6] using Bayesian Binary Method (BBM) in Reconstruct-Ancestral-States-in-Phylogenies 4.0 (RASP4) [25, 26]. The ML phylogeny obtained with IQ-TREE was used for the BBM analysis, which was conducted as two independent runs of 10 chains that ran for 6,000,000 generations and were sampled every 100 generations. The fixed Jukes-Cantor+Gamma evolutionary model was implemented for each run. DnaSP was used to calculate various statistics of genetic variation, including haplotype diversity, nucleotide diversity, theta based on the number of segregating sites, and number of polymorphic sites for separate mtDNA control region *Callithrix* clades. *Callithrix* clades were those based on the results of the phylogenetic analyses. A NeighborNet network of mtDNA haplotypes of all sampled control region sequences was made with default settings in SplitsTree4 [27].

### 2.4 Mitogenome Alignment and Data Analysis

For mitogenome analysis, we specifically focused on *C. jacchus* mitogenomes, but used publicly available *Cebuella pygmaea* (GenBank accession# MZ747454.1), *C. penicillata* (GenBank accession# NC_030788.1), *C. geoffroyi* (GenBank accession# LR745201), and *C. kuhlii* (GenBank accession# KR817257) as phylogenetic outgroups. All mitogenomes were initially aligned in MAFFT with default settings and MAFFT alignments were confirmed visually in Mesquite. The alignment was 16554 bases. Unique mitogenomic haplotypes were determined with DnaSP and then used for ML phylogenetic reconstruction with IQ-TREE and Bayesian phylogenetic reconstruction with MrBayes. For the ML phylogeny, we used the optimal substitution model TIM2+F+I+G as calculated with ModelFinder in IQ-TREE under BIC, kl. We performed the ML analysis two separate times in IQ-TREE with 1000 ultrafast bootstrap (BS) replications per run. The two resulting ML trees were then compared visually for concordance. For the Bayesian tree, we used Modeltest with the -T flag set to “mrbayes” to determine the optimal substitution model as “HKY+I+G4” with BIC. The Bayesian tree was reconstructed via Markov Chain Monte Carlo (MCMC) runs with 1,000,000 generations and tree and parameter sampling occurring every 100 generations. Upon completion of the two runs, the first 25% of generations were discarded as burn-in. To check convergence of all parameters and the adequacy of the burn-in, we assessed PSRF and that all ESS values were above 200. Phylogenetic trees were visualized and edited with FigTree. We made a MAFFT alignment of *C. jacchus* only mitogenomic sequences which was then used to make a NeighborNet haplotype network with default settings in SplitsTree4. DnaSP was used to calculate various statistics of genetic variation, including haplotype diversity, nucleotide diversity, theta based on the number of segregating sites, and number of polymorphic sites for the *C. jacchus* mitogenomic clade. Due to the lack of known provenance for many of the utilized *Callithrix* mitogenomes, we opted to not perform mitogenomic biogeographic analysis for this data set.

## 3. Results

### 3.1 *Callithrix* Mitochondrial Phylogenetic Analysis

For the mtDNA control region data, branches of the Bayesian and ML phylogenetic trees (Figures 2A-B), respectively, possessed overall strong statistical support with the exception of some weak support towards the tree tips. MtDNA control region haplotypes clustered into eight main species-level clades, with *C. aurita* being the most basal clade. The *C. penicillata* and *C. kuhlii* clades were polyphyletic, with the former forming three clades and the latter forming two clades. Within the *C. penicillata* clades, haplotypes with known provenance clustered together according to similar geographical origin. It was not clear if this is also the case for haplotypes with the *C. kuhlii* clades as their provenance was unknown. *Callithrix jacchus* haplotypes formed the largest and one of the most recent clades of the two phylogenies. Haplotypes from known hybrids clustered within various clades in both phylogenies. There were two instances of discordance between an individual’s phenotype and mtDNA control region haplotype within the *C. penicillata* Caatinga clade, (BJT100 haplotype/ *C. jacchus* phenotype; BJT22 haplotype/*C. aurita* phenotype). There was also phenotype-haplotype discordance within the *C. jacchus* clade (BJT020 haplotype/*C. aurita* phenotype, BJT79 haplotype/*C. aurita* phenotype, CPE021 haplotype/*C. penicillata* phenotype, and CGE001 haplotype/*C. geoffroyi* phenotype).

**Figure 2:**
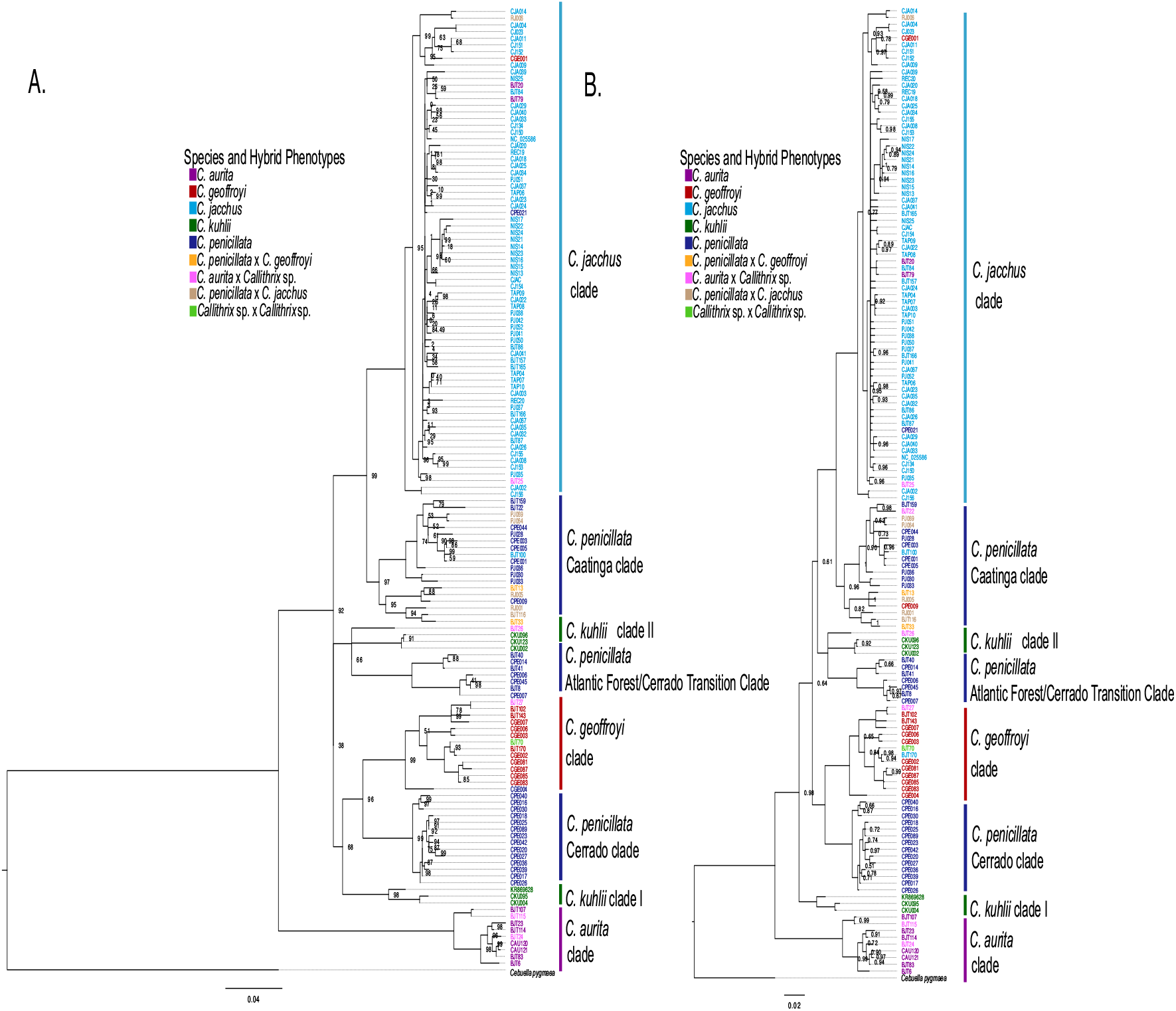
Phylogenetic relationships among *Callithrix* mtDNA control region. (A) Maximum likelihood phylogenetic relationships among haplotypes for which node support is shown where bootstrap support <70% in the tree. (B) Bayesian phylogenetic tree for which node support is shown where bootstrap support <1.0 in the Bayesian tree. In both plots, newly sequenced mtDNA control region sequences are showed in bold and are italicized. Haplotype colors at tips correspond to the ‘Species and Hybrid Phenotypes’ legend, and indicate phenotypes associated with each given haplotype.

The control region ML and Bayesian phylogenies show similar topologies except in respect to the *C. jacchus* clade. The Bayesian phylogeny was not able to resolve relationships between *C. jacchus* clade haplotypes. The ML tree showed better resolution, but lacks strong statistical branch support within the *C. jacchus* clade. The ML phylogeny did possess one well-supported *C. jacchus* subclade of haplotypes originating from the Nísia Floresta Field Station in Pernambuco state (NISXXX haplotypes in Figure 3). Newly sequenced control region haplotypes from international captive facilities all grouped within the *C. jacchus* clade, thus showing concordance with phenotypes of sampled animals. The included CLEA marmoset control region sequences all shared haplotype CJ154 and those from Company A all shared haplotype CJ134. On the other hand, marmoset control region haplotypes from NEPRC formed two separate clusters, one composed of CJ153 and CJ155 and the other composed of CJ151 and CJ152. No control region haplotypes from across the three facilities clustered together.

**Figure 3:**
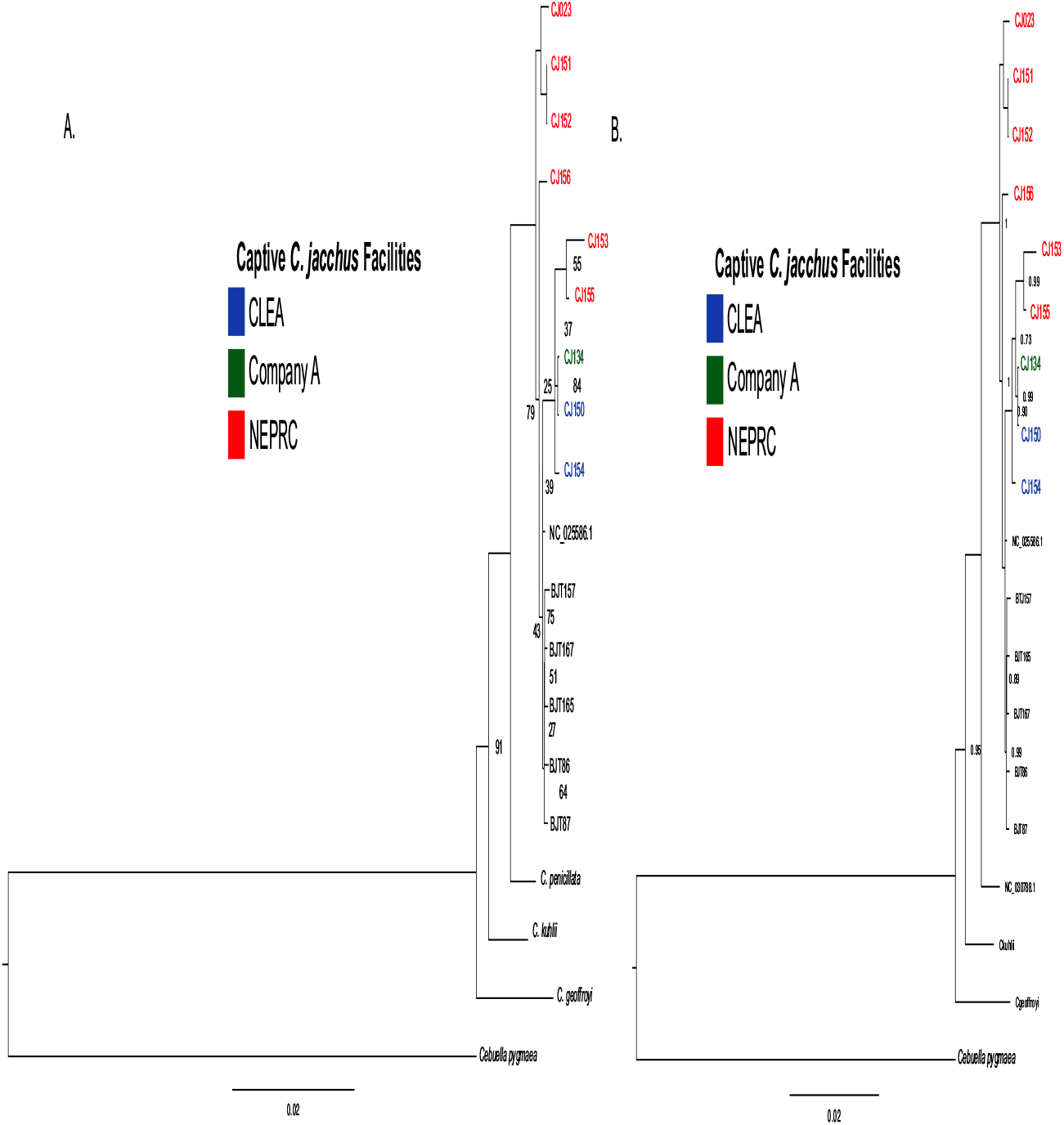
Phylogenetic relationships among mitogenomic haplotypes of *C. jacchus* lineage with focus on marmosets from captive research facilities. (A) Maximum likelihood phylogenetic relationships haplotypes for which node support is shown where bootstrap support <70% in the tree. (B) Bayesian analysis of mitogenomes for which node support is shown where bootstrap support <1.0 in the Bayesian tree. Newly sequenced mitogenome haplotypes are colored. These haplotype colors at tips correspond to the ‘Captive *C. jacchus* Facilities’ legend, and indicate the captive facility from where a given haplotype originates.

The mitogenomic Bayesian and ML trees (Figure 3A-B) show stronger phylogenetic resolution and statistical support among lineages of the *C. jacchus* clade relative to the mtDNA control region phylogenies. Mitogenome haplotypes (BJTXXX) included from marmosets of known provenance within the Caatinga formed their own well supported subgroup within the *C. jacchus* clade. Newly sequenced mitogenomes from marmosets samples from international captive facilities show similar phylogenetic clustering patterns as that for the mtDNA control region, but with better resolution of divergence patterns among mitogenomic haplotypes. In the case of mitogenomes included from CLEA marmosets, we see the lineages collapse into two separate haplotypes. Also, none of the captive facility haplotypes clustered within the Caatinga *C. jacchus* mitogenomic lineages.

### 3.2. *Callithrix* Mitochondrial Network Analyses

Due to poor phylogenetic resolution of relationships of haplotypes within the *C. jacchus* mtDNA control region clade, we constructed a NeighborNet network (Figure 4A) to try to parse out these relationships. This network resulted in a star-like pattern, which further indicates that the *C. jacchus* clade represents relatively recently *Callithrix* mtDNA lineages. The Nísia Floresta subclade is also present within the network towards the upper right side. There seems to exist another distinct subclade that stands out on the left side of Figure 4A that includes haplotypes CJ156, CJA002, CGE001, etc. The original provenance of haplotypes forming this second subclade is unknown, so it is not possible to assign them to a possible geographic origin.

**Figure 4:**
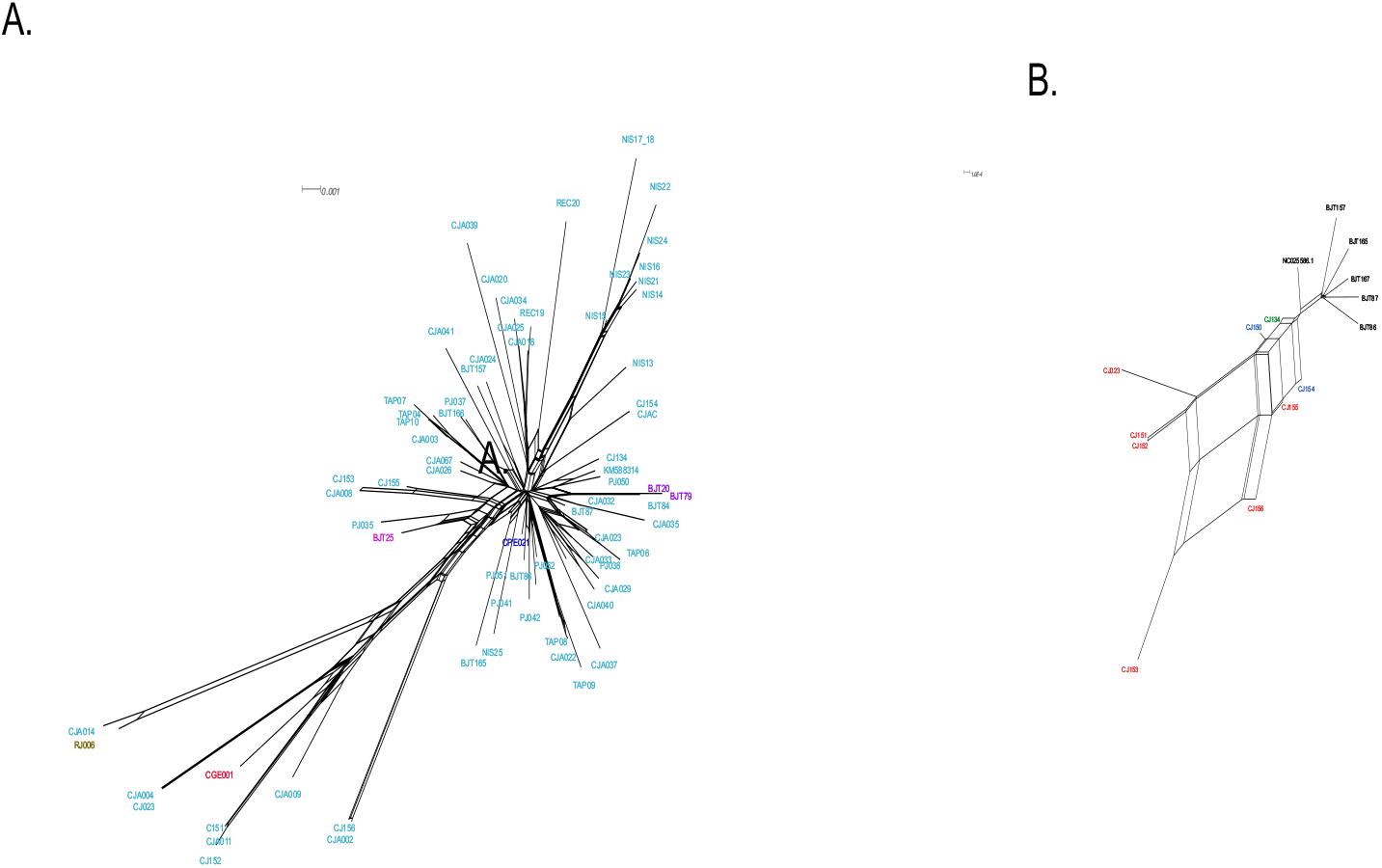
Neighbor network of haplotypes from phylogenetic *C. jacchus* clade. Haplotype colors at tips follow Fig. 2 ‘Species and Hybrid Phenotypes’ legend, and indicate phenotypes associated with each given haplotype.

In comparison to the mtDNA control region, the *C. jacchus* mitogenome NeighborNet network (Figure 4B) shows relatively better haplotype resolution. The mitogenome network shows a distinct clade for the BJT Caatinga haplotypes. Haplotypes from NEPRC (CJ023, CJ151, CJ152, CJ153, CJ155, and CJ156) cluster similarly as in the phylogenetic trees. However the network suggests a closer relationship among these haplotypes than in the phylogentic trees. There were two distinct mitogenomic haplotypes from CLEA marmosets (CJ150 and CJ154), which did not cluster closely.

### 3.3. MtDNA Diversity of *Callithrix* Taxa

Calculations of *Callithrix* diversity of the mtDNA control region were performed for the major clades defined by the above phylogenetic analysis (Table 2). We made an exception for the two *C. kuhlii* clades, which we collapsed together due to low intraclade sample numbers. Haplotypes from hybrids and haplotypes that were incongruent with individual phenotypes were excluded from the diversity analysis. The *C. jacchus* clade possessed the highest number of haplotypes, haplotype diversity, and number of segregating sites. However, despite highly variable sampling numbers for each clade, we see similar values of inter-clade haplotype diversity. For nucleotide diversity, we see the highest values in the collapsed *C. kuhlii* clade followed by the *C. geoffroyi* clade, and the lowest value in the *C. penicillata* Cerrado clade. We find similar patterns among these three clades for estimates of per sequence theta, but with the positions of *C. kuhlii* and *C. geoffroyi* clades being reversed. The *C. aurita* clade maintains an intermediate position across all diversity measures.

**Table 2.** Summary of *Callithrix* MtDNA Control Region Diversity Statistics

| Taxon | Sequences | Total<br>Number<br>of<br>Sites | Haplotype<br>Number | Haplotype<br>Diversity | Nucleotide<br>Diversity | Theta<br>(per<br>sample) | Polymorphic<br>Sites |
| --- | --- | --- | --- | --- | --- | --- | --- |
| *C. aurita* | 9 | 782 | 7 | 0.944 | 0.021 | 15.82 | 43 |
| *C. geoffroyi* | 16 | 789 | 12 | 0.958 | 0.032 | 27.42 | 91 |
| *C. jacchus* | 102 | 791 | 62 | 0.981 | 0.028 | 22.51 | 117 |
| *C. kuhlii* | 9 | 796 | 6 | 0.889 | 0.043 | 25.02 | 68 |
| *C. penicillata*<br>Caatinga Clade | 18 | 797 | 10 | 0.935 | 0.029 | 21.51 | 74 |
| *C. penicillata*<br>Cerrado Clade | 24 | 795 | 13 | 0.924 | 0.11 | 8.84 | 33 |
| *C. penicillata*<br>Cerrado-Atlantic<br>Forest Clade | 11 | 795 | 7 | 0.927 | 0.023 | 13.31 | 39 |

**Table 3.** Summary *C. jacchus* Mitogenome Diversity Statistics

| Taxon | Sequences | Total<br>Number<br>of<br>Sites | Haplotype<br>Number | Haplotype<br>Diversity | Nucleotide<br>Diversity | Theta<br>(per<br>sample) | Polymorphic<br>Sites |
| --- | --- | --- | --- | --- | --- | --- | --- |
| *C. jacchus* | 1 | 16262 | 9 | 0.892 | 0.002 | 27.73 | 92 |

### 3.4. Biogeography of *Callithrix* mtDNA control region clades

The ancestral origins of *Callithrix* phylogenetic mtDNA control region clades and subclades based on BMM biogeographic analysis were mostly concordant with assigned origins of mtDNA haplotypes (Figure 5). Within the *C. aurita* clade, haplotypes from node 284 in Figure 5 originate in the state of Rio de Janeiro, as expected. However, for the remaining *C. aurita* clade haplotypes, BMM analysis was not able to determine more precise biogeographic origins of haplotypes beyond the general known geographic distribution of *C. aurita*. For the two *C. kuhlii* clades, BMM analysis placed both clades within the Atlantic Forest of Bahia state. While all *C. kuhlii* haplotypes were of unknown provenance, their BMM biogeographic designations are in line with the known distribution of *C. kuhlii*. Among haplotypes of the *C. geoffroy*i clade, while BMM analysis placed all haplotypes into the known geographic range of *C. geoffroyi* with high posterior probability (PP), the analysis did not show geographic structuring within the clade. However, haplotypes were included with known provenance from the Caatinga biome of Minas Gerais state (BJT143) and Atlantic Forest of Espírito Santo state (BJT 170). These particular haplotypes cluster into two separate subclades that diverge at node 247. BMM biogeographic origins of the three *C. penicillata* clades also received strong PP estimates that were concordant with geographic separation of the clades, as was already apparent in the phylogenetic analysis.

**Figure 5:**
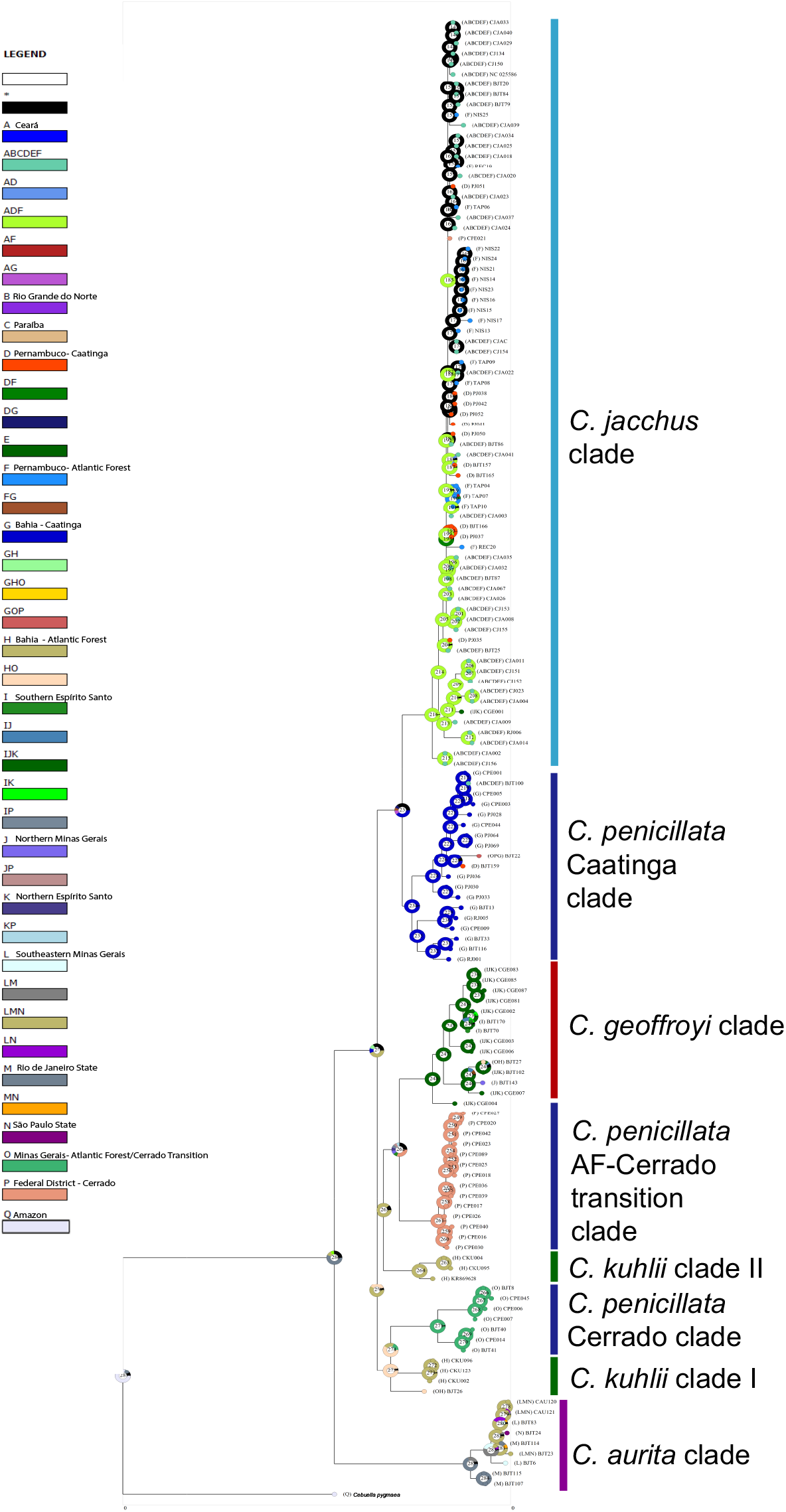
Ancestral state reconstructions performed by the Bayesian Binary MCMC analysis as implemented in RASP using the ML rooted tree. Donut charts at each node represent ancestral host estimations in terms of posterior probability (PP). Each node is internally identified with a number. Localities where species associated with each phylogenetic clade were sampled or known to occur are color coded in the map inset on the left side.

Within the *C. jacchus* clade, BMM analysis placed the nodes of the bottom half of the clade within a combined biogeographic region of the states of Ceará and Pernambuco with high PP. Yet, the biogeography of nodes falling within the upper half of the *C. jacchus* clade could not to be determined by BMM analysis. Nodes within the *C. jacchus* clade for which BMM determination agreed with known haplotype provenance included the Atlantic Forest of Pernambuco state at node 189 and 190 and the Caatinga of Pernambuco state for PJ037 and BJT166 at node 193. We were not able to resolve the probable biogeographic origins of newly sequenced control region sequences from *C. jacchus* haplotypes obtained from international captive facilities.

## 4. Discussion

With the exception of a single haplotype (BJT100), all haplotypes from marmosets with a *C. jacchus* phenotype, including those sampled at international facilities, clustered within the *C. jacchus* phylogenetic clade. All phylogenies possessed relatively shallow branches within the *C. jacchus* clade compared to the rest of *Callithrix* clades. Previous divergence dating of the *Callithrix* mitogenomic phylogeny shows that the *C. jacchus* clade arose approximately 0.5 million years ago (MYA) [6]. Thus, *C. jacchus* clade haplotypes from mtDNA are among the youngest within the *Callithrix* genus, which certainly accounts for the short branch lengths within this clade. Overall, mitogenomic phylogenies provided relatively better resolution of *C. jacchus* haplotype divergence than that of the control region. There were however slight differences in *C. jacchus* haplotype branching topology for the two mitogenome phylogenies. The Bayesian mitogenomic tree showed better support for *C. jacchus* branching patterns than the ML mitogenomic tree.

Due to the shallow branches within the *C. jacchus* mitogenomic and mtDNA con-trol regions clades, networks of *C. jacchus* haplotypes were constructed to complement phylogenetic approaches. The star-like pattern of *C. jacchus* clade mtDNA control region networks produced in our study as well as that of [4] provide further evidence of the relatively young age of this clade. The mitogenomic *C. jacchus* clade network showed somewhat clearer resolution of network paths among haplotypes. Haplotypes from marmosets originating from the Caatinga biome (those whose names begin with BJT in Figure 4B) form a clear group that is exclusive to the other haplotypes. We also see this grouping within the mitogenomic phylogenies, but neither approach can further resolve relationships among these haplotypes.

*Callithrix jacchus* genetic diversity based on the mtDNA control region was intermediate relative to other *Callithrix* species. Genetic diversity measures estimated for the mitogenome for *C. jacchus* were similar to that of the mtDNA control region in terms of haplotype diversity but an order of magnitude smaller in terms of nucleotide diversity. Based on differences in haplotype numbers observed for different biomedical facilities included in this study (see Figure 3), it is likely that levels of mitochondrial genetic diversity differ among captive *Callithrix* colonies. Among the international facilities from which we sampled marmosets, it is likely that NEPRC possesses the highest relative level of mtDNA genetic diversity given that marmosets from this facility possessed the highest number of mtDNA and mitogenome haplotypes. It is likely that the CLEA colony would have possessed the lowest relative level of mtDNA genetic diversity given that marmosets from this facility possessed a single haplotype for both mitochondrial datasets. To fully assess the genetic diversity within and among international biomedical facilities, a larger number of animals will need to be sampled and analyzed.

### 4.1 Biogeography of *C. jacchus* mtDNA Control Region Haplotypes

Unlike other *Callithrix* phylogenetic clades, assignment of biogeographic origin of *C. jacchus* control region haplotypes was not fully successful over the entire clade. Haplotypes from *C. jacchus* in biomedical facilities were assigned to a biogeographic origin within the Atlantic Forest, but a more specific location of origin was not possible. The biogeographic origins for a number of other *C. jacchus* haplotypes could not be resolved, even when they were associated with marmosets of known provenance. Haplotypes with known provenance whose biogeographic origin was correctly assigned included PJ037 and BJT166 which originate from the Caatinga and TAP10, TAP07, and TAP04 which originated from the Atlantic Forest of Pernambuco state. Previously published *Callithrix* biogeographic analysis based on the complete mitogenome performed better for the *C. jacchus* clade, as biogeographic origins were assigned for all haplotypes [6].

Due to the higher phylogenetic and biogeographic resolution of *Callthrix* haplotypes provided by the mitogenome in comparison to the mtDNA control region, the *C. jacchus* biogeographic results presented here should be interpreted with caution. An increased availability of known-provenance mitogenomes from across the *Callithrix* range would certainly improve the precision of *Callithrix* biogeographic analysis. In the specific case of marmosets from biomedical facilities, we are missing the necessary genetic and geographic information from the natural *C. jacchus* range to more precisely determine their biogeographic origins. Once such information becomes more accessible, based on the previously published results discussed above [6], we suggest that data from the entire mitogenome will provide greater resolution for biogeographic analysis than data from the control region alone.

Based on several published mtDNA phylogenies [4, 6, 9], as well as those presented here, one plausible biogeographic hypothesis is that the *C. jacchus* clade originated in the Caatinga. Although *C. jacchus* occurs naturally in the Caatinga, Cerrado, and northeastern Brazilian Atlantic Forest biomes (Figure 1), mitochondrial phylogenies show that the species diverged from the *C. penicillata* Caatinga clade. Further, biogeographic modeling suggests that the *C. jacchus/C. penicillata* ancestor dispersed into the Cerrado, then *C. penicillata* became isolated in either the Cerrado or Caatinga and subsequently *C. jacchus* dispersed across Caatinga, Cerrado, and the Atlantic Forest [28]. It is plausible then that an ancestral *C. penicillata* population may have dispersed from the Cerrado into the Caatinga, and later diverged into the *C. jacchus* and *C. penicillata* Caatinga clade.

A possible Caatinga origin for *C. jacchus* carries several implications for the evolutionary biology and biomedical use of this species. Stark environmental differences between the humid Atlantic Forest and semiarid Caatinga and Cerrado biomes may have driven genetic, phenotoypic, and behavioral differences in *C. jacchus* subpopulations found within these areas (personal obs., Malukiewicz). Reduced or absent gene flow between subpopulations spread across a wide geographic range can result in notable genetic differences within the same species [29]. Whether any such genetic differences exist among natural *C. jacchus* subpopulations is currently unknown. We also do not know how any existing genetic and adaptive differences between native *C. jacchus* populations have played into the population genetics of *C. jacchus* in biomedical colonies outside of Brazil. Resolving these questions is important for optimizing biomedical use of marmosets, particularly as genetic differences associated with ancestry and geographic origin of non-human primates can influence results of biomedical experiments dependent on non-human primate models [29–32].

### 4.2 Hybridization within the C*allithrix* Genus based on Mitogenomic and Mitochondrial Markers

The control region phylogenies showed a number of haplotypes within the *C. jacchus* clade that are discordant with the phenotypes of sampled marmosets (BJT20, BJT79, CPE021, CGE00). This same pattern has been previously observed for BJT20 and BJT79 in a mitogenomic phylogeny and attributed to genetic introgression of *C. jacchus* mtDNA into the genetic background of *C. aurita* as a result of anthropogenic hybridization [6]. In this study and a previous one [4], haplotypes CPE021 and CGE001 grouped within *C. jacchus* mtDNA clades, but were derived with individuals sampled from captive facilities in the US with *C. penicillata* and *C. geoffroyi* phenotypes, respectively. These haplotypes may represent cases of incomplete lineage sorting if the ancestors of the sampled marmosets never interbred with *C. jacchus*, as incomplete lineage sorting should not promote geographical proximity of interspecifically shared alleles under local introgression [33].

The case of haplotype BJT100 represents a cryptic hybrid, and was associated with an individual sampled in a captive facility in Petrolina, Pernambuco, Brazil. This facility is located at the natural border between the natural geographic ranges of *C. jacchus* and *C. penicillata* where a natural hybrid zone exists [4]. However, the two species are also frequently captured illegally, interbred, and kept as pets in Brazil [1, 5].

In many cases hybridization seems to reduce fitness globally, and at the cellular level hybrids may face challenges in reconciling protein interactions [34, 35]. Our current results for marmosets from international biomedical facilities do not indicate the presence of hybrids among our sampled individuals. However, there is phenotypic evidence of possible *Callithrix jacchus* x *C. penicillata* hybrids present among marmosets kept in US biomedical facilities (personal observation, Malukiewicz). Additionally, previous mtDNA genetic studies show that cryptic hybridization in the wild does seem to occur regularly within the Callithrix genus [4–6, 9].

Accordingly, we would expect that if hybrid marmosets are present in US biomedical facilities, they may not may not show obvious phenotypic signs of interspecific ancestry. Thus, we do not recommend relying solely on phenotype for species identification of marmosets maintained in captivity. It will also be valuable to perform mitogenomic analysis of captive animals as additional support for their ancestry. Eventually, genome-wide autosomal variants will be identified that can provide a more complete picture of ancestry than mitogenomic analysis alone. Such checks could help minimize spurious results in future research looking to utilize marmosets as biomedical models.

## 5. Conclusions

The mitogenomic network and phylogenetic results from this and other work [6, 9, 10] show that analyses of the mitogenomic variation holds substantial potential for resolving evolutionary relationships among *C. jacchus* populations, defining species boundaries, assigning individuals to populations, and detecting hybrids. The mitogenome is an especially useful genetic marker for non-model taxa, taxa which are presently deficient in genetic data, or little-studied taxa, as it can be used relatively quickly to access ancestry particularly using approaches such as genomic skimming [9]. Within the US, significant effort and funding are being directed toward assessing whole genome genetic diversity and genotyping among marmosets at biomedical facilities motivated by increasing interest in marmosets as biomedical model for neurodegenerative and other diseases [1]. Mitogenomes can be mined from whole genome sequencing data [6, 9]. Thus, such mined data can be obtained from marmosets at international captive facilities as whole genomic data becomes increasingly available from these primates. If sampling and genomic sequencing efforts can be also matched for natural *C. jacchus* populations in Brazil, utilization of mitogenomic data from wild *C. jacchus* of known provenance can be harnessed to determine the biogeographic origins of captive *C. jacchus*. Expanding our efforts into understanding the evolutionary history of marmosets will greatly enhancethe use of marmosets for biomedical purposes. Thus, combining marmoset mitogenomic data with phenotypic and nuclear genomic data is crucial for the management of captive marmoset populations with the goal of maintaining genetic diversity, organizing appropriate breeding plans, identifying species and hybrids, and selecting animals with appropriate genetic backgrounds for biomedical research.

## Acknowledgements

This work was supported by the Stanley Center for Psychiatric Research at the Broad Institute of MIT and Harvard, the James and Patricia Poitras Center for Psychiatric Disorders Research at MIT and the Hock E. Tan and K. Lisa Yang Center for Autism Research at MIT. This work was supported by DAAD doctoral fellowship to JLGF, a Brazilian CNPq Jovens Talentos Postdoctoral Fellowship (302044/2014-0), a Brazilian CNPq DCR grant (300264/2018-6), a Marie-Curie Individual Fellowship (AMD-793641-4), an American Society of Primatologists Conservation Small Grant, and an International Primatological Society Research Grant for JM. We would like to thank Dr. Anne Stone for her methodological assistance and input and Dr. Carlos Ruiz-Miranda for assistance with samples. We thank Vanner Boere, Ita de Oliveira, Daniel S. Silva, the Guarulhos Zoo, the CEMAFAUNA, the CPRJ staff, SERCAS staff, and AMLD staff for assistance with wild and captive populations.

